# An open state of a voltage-gated sodium channel involving a π-helix and conserved pore-facing Asparagine

**DOI:** 10.1101/2021.07.28.454140

**Authors:** Koushik Choudhury, Marina A. Kasimova, Sarah McComas, Rebecca J Howard, Lucie Delemotte

## Abstract

Voltage-gated sodium (Nav) channels play critical roles in propagating action potentials and otherwise manipulating ionic gradients in excitable cells. These channels open in response to membrane depolarization, selectively permeating sodium ions until rapidly inactivating. Structural characterization of the gating cycle in this channel family has proved challenging, particularly due to the transient nature of the open state. A structure from the bacterium *Magnetococcus marinus* Nav (NavMs) was initially proposed to be open, based on its pore diameter and voltage-sensor conformation. However, the functional annotation of this model, and the structural details of the open state, remain disputed. In this work, we used molecular modeling and simulations to test possible open-state models of NavMs. The full-length experimental structure, termed here the α-model, was consistently dehydrated at the activation gate, indicating an inability to conduct ions. Based on a spontaneous transition observed in extended simulations, and sequence/structure comparison to other Nav channels, we built an alternative π-model featuring a helix transition and the rotation of a conserved asparagine residue into the activation gate. Pore hydration, ion permeation and state-dependent drug binding in this model were consistent with an open functional state. This work thus offers both a functional annotation of the full-length NavMS structure, and a detailed model for a stable Nav open state, with potential conservation in diverse ion-channel families.

## Introduction

Voltage gated sodium (Nav) channels are membrane proteins that play an important role in the propagation of action potentials in excitable cells during nerve impulse conduction, among other physiological processes. These channels are involved in cardiac, muscular and neurological disorders, making it important to understand the mechanisms that underlie their function (1). When the membrane reaches a threshold potential in a prototypical nerve cell, Nav channels open to allow Na^+^ ions to flow inward, down their electrochemical gradient. Key to their function is a subsequent rapid inactivation which stops the ion flow, and leaves time for the slow voltage-gated potassium channels to open, letting K^+^ ions out and ultimately returning the cell to its resting potential (2).

An eukaryotic Nav channel comprises a single 2000-residue polypeptide chain, with four homologous domains arranged in a pseudotetrameric architecture, as verified by a handful of recent cryoEM structures reported in recent years (3–8). In contrast, bacterial Nav channels are homotetramers with ∼270 residues per subunit and have a simpler architecture with smaller intracellular and extracellular domains (9–26). Despite their limited (∼25%) sequence identity, bacterial Nav channels have been shown to share drug sensitivity and other functional properties with their eukaryotic counterparts, making them compelling model systems for structure/function studies (27).

Bacterial Nav channels consist of three main domains - a voltage sensing domain (VSD), a pore domain and a C-terminal domain (Figure 1.A) (28). Each subunit VSD consists of four helices labelled S1-S4, arranged in a bundle. S4 contains several basic residues, which are positively charged at neutral pH and are responsible for sensing the changes in potential difference across the membrane. The VSDs are assigned to an activated conformation when these arginines are displaced towards the extracellular side, relative to the resting conformation (10). The pore domain consists of the tetrameric arrangement of the S5 and S6 helices from each of the four subunits. The primary sequences of the pore domain and VSD are connected by a ∼20 residue-long S4-S5 linker, which connects the pore domain of one subunit to the VSD of another subunit, in a so-called domain-swapped arrangement. The C-terminal domain includes a partially disordered neck region, and a coiled coil at the C-terminus of the protein. In the resting state of the channel, the VSDs are presumed to occupy a resting conformation, and the pore is closed. In the open state, the VSDs are activated, and the pore is open. In the inactivated state, the VSDs remain activated, but the pore no longer allows ions to pass. The role of the C-terminal domain in the functional cycle is not yet fully established.

**Figure 1:**
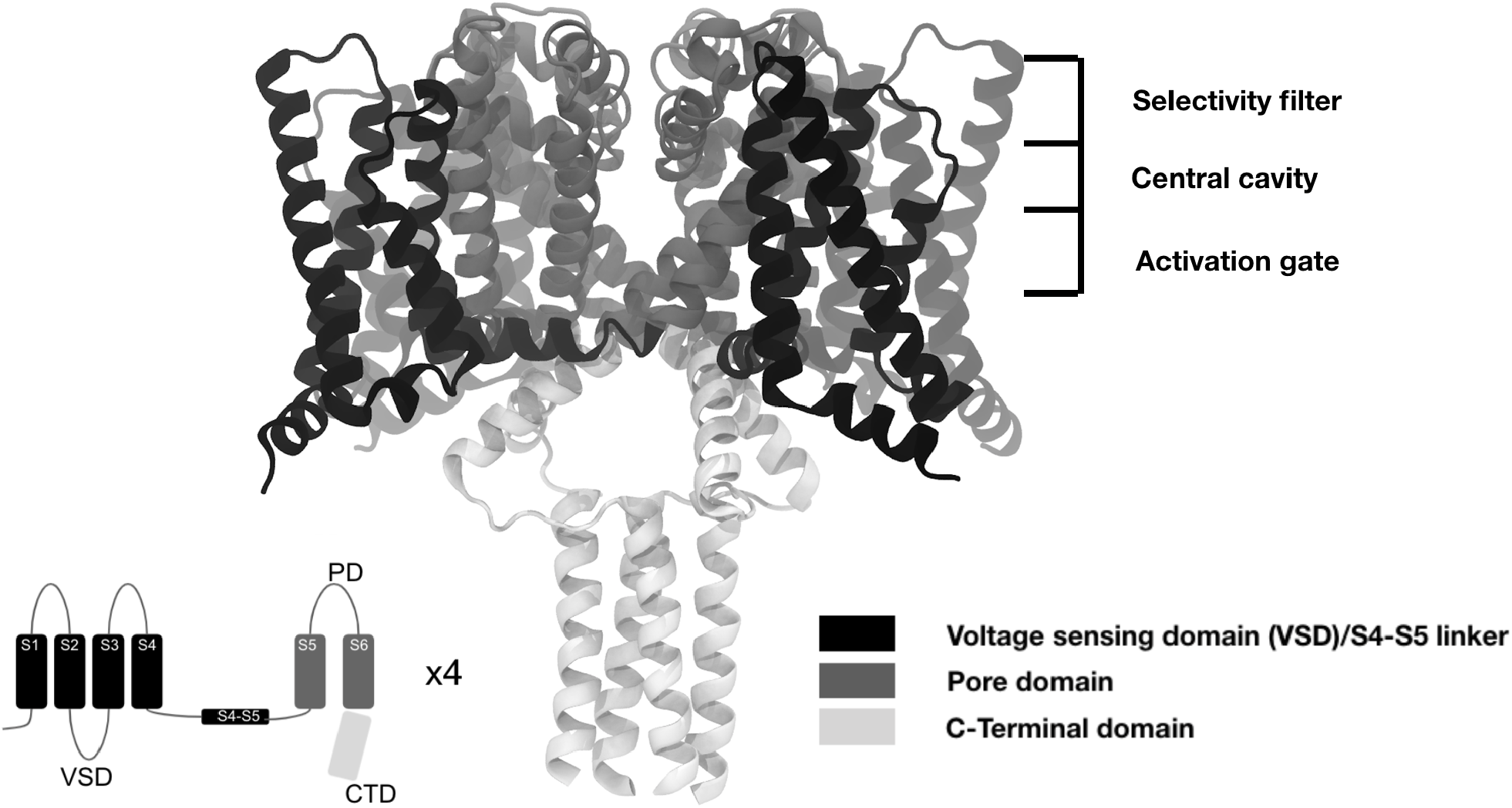
Ribbon representation of the full-length NavMs X-ray structure (PDB ID 5HVX) with the voltage-sensor domain (VSD), pore domain, and C-terminal domain represented in dark, medium, and light gray, respectively. Insert shows a cartoon of a single NavMs subunit, colored as in the 3D model.

Many efforts have been made to determine the structure of Nav channels in different functional states. Bacterial Nav channels have provided some insights, including structures of at least six subtypes (NavAb, NavMs, NavRh, NavCt, NaChBac and NavAe) in apparently distinct states (9–26). Still, these structures were solved in a non-native environment (at cryogenic temperatures, solubilized by detergents) and often in the presence of mutations, so their functional assignments can be ambiguous.

The resolution of open-state structures has proved particularly challenging. Because this state transitions spontaneously to the inactivated state under physiological conditions, trapping it for timescales sufficient for structure determination has often made it necessary to resort to protein modifications. In the case of NavMs, a bacterial Nav from *Magnetococcus marinus*, a full-length channel structure thought to be trapped in a conductive state was determined, based on the radius of the pore (24). However, several other factors, besides the pore radius, dictate whether a pore is conductive, such as hydrophobicity, hydration, and interaction with ions (29–33). In addition, the open state should enable access of open-pore blocker drugs to their binding site via a hydrated pathway across the gate (34, 35).

Molecular dynamics (MD) simulations enable the direct visualization of atomistic interactions in and around a channel pore, including water, ions, or drugs, and can thus contribute to the functional assignment of experimental structures (30). MD simulations of the full-length NavMs structure have indeed questioned its functional assignment as an open state: its pore gate tended to dehydrate, even in the presence of stabilizing interactions with the other channel domains (36). Thus the functional annotation of this structure, and its relationship, if any, to a stable open state, remain unclear.

In this work, we used molecular modeling and MD simulations to test the open-state properties of NavMs models. Consistent with previous indications, the full-length X-ray structure was dehydrated; furthermore, even supraphysiological electric fields did not trigger wetting of the activation gate. Based on spontaneous transitions observed in these simulations, and sequence and structural features of related channels, we then constructed an alternative open-state model by introducing a π-helix in S6, N-terminal to the activation gate. Hydration, ion permeation, and drug binding in this model supported its annotation as a putative open state, providing a newly detailed testable mechanism for gating in the Nav family.

## Materials and Methods

### Model and simulation system building

Simulations in this work were based on the full-length X-ray structure of NavMs (PDB ID 5HVX), and incomplete loops built using MODELLER 9.22 (37). The π-model was built by inserting an unpaired backbone hydrogen bond in S6 using MODELLER. In aligning the template and target sequences, one gap was introduced in the template immediately prior to Thr-207, and a second in the target following Thr-234, thus shifting the target sequence between Thr-207 and Thr-234 upstream by one position (Figure S1). This location of the insertion mimics the π-helix transition observed in a related TRP channel (38). Mutations were inserted using Visual Molecular Dynamics (VMD) (39). Each model was embedded in a homogenous lipid bilayer consisting of 362 to 400 1-palmitoyl-2-oleoyl-*sn*-glycero-3-phosphocholine (POPC) molecules using the CHARMM-GUI Membrane builder (40). The system was hydrated by adding a ∼45-Å layer of water to each side of the membrane. Lastly, the system was ionized to reach a 150-mM NaCl concentration. Complexes with lidocaine or flecainide were prepared by randomly placing the drug molecule in the central cavity, and building the remainder of the system as described above. The Charmm36 force field was used to describe interactions between protein (41), lipids (42) and ions. The TIP3P model was used to describe the water particles (43).

### Drug molecules parametrization

The parameters for lidocaine and flecainide were generated using CGENFF (44). Lidocaine has a pKa of 7.56, thus it is 70% charged and 30 % neutral at physiological pH. Flecainide has a pKa of 9.3, thus it is 99% charged and 1% neutral at physiological pH (45). The charged forms of these drugs block the pore by entering it from the intracellular side via the so-called hydrophilic pathway (46–48). To evaluate the propensity of open pore models to allow the binding of open pore binders, we thus chose to model the charged form of lidocaine and flecainide. The lidocaine parameters were checked by calculating the free energy for water to membrane transitions (Figure S2). These free energies were estimated using the accelerated weight histogram (AWH) method in GROMACS, simulating 6 walkers for 200 ns each. The center of mass between a sodium ion restrained to its initial position and lidocaine was chosen as the collective variable. The free energy profile and the preferential drug localization below the lipid/solution interface agrees with the results from previous studies (49). In addition, the drug-membrane partition coefficient for charged lidocaine (log(P)) is 1.49 (50), yielding a free energy of lipid/water partitioning ΔG ∼ -8.5 kJ/mol, in qualitative agreement with the free-energy difference determined in our simulations (ΔG ∼ -11 kJ/mol). Although experimental partition coefficients were not readily available for charged flecainide, based on this validation with lidocaine, we proceeded with CGENFF parameters for both drugs.

### Simulation parameters

The systems were minimized for 15000 steps using steepest descent, and equilibrated with a constant number of particles, pressure and temperature (NPT) for 30 to 45 ns, during which the position restraints were gradually released. During equilibration, pressure was maintained at 1 bar through Berendsen pressure coupling; temperature was maintained at 300 K through Berendsen temperature coupling (51) with the protein, membrane and solvent coupled. Finally, unrestrained production simulations were run, using a timestep of 2 fs, and Parinello-Rahman pressure coupling (52) and Nose-Hoover temperature coupling (53). The LINCs algorithm (54) was used to constrain bonds involving hydrogen atoms. For long-range interactions, periodic boundary conditions and particle mesh Ewald (PME) were used (55). For short-range interactions, a cut-off of 12 Å was used. Simulations were performed using GROMACS (versions 2016, 2019 and 2020) (56, 57). A transmembrane voltage of +750 mV or -750 mV was applied in the form of an external electric field to the equilibrated NavMs α-model to study electrowetting. The transmembrane voltage was calculated as the product of the electric field by the simulation box length along the membrane normal. Equilibrium simulations described in this work are summarized in Table 1.

**Table 1:** Summary table of equilibrium simulations performed herein.

| System | Equilibration | Production |
| --- | --- | --- |
| NavMs α-model | 48 ns | 400 ns |
| NavMs α-model +750mV | start from end of NavMs α-model production | 800 ns |
| NavMs α-model -750mV | start from end of NavMs α-model production | 8000 ns |
| NavMs π-model | 48 ns | 800 ns |
| NavMs α-model - Lidocaine | 36 ns | 200 ns |
| NavMs π-model-Lidocaine | 36 ns | 200 ns |
| NavMs α-model - Flecainide | 36 ns | 200 ns |
| NavMs π-model - Flecainide | 36 ns | 200 ns |
| NavMs π-model N212A | 48 ns | 505 ns |
| NavMs π-model N212L | 48 ns | 452 ns |
| NavMs π-model N212C | 48 ns | 456 ns |

### Ion permeation free energy calculation

The free energy of permeation of a Na^+^ ion along the channel pore axis was calculated using the accelerated weight histogram (AWH) method (58). For each equilibrated structure, we applied an independent AWH bias and simulated 6 walkers for 200 ns each (representing a total of 1200 ns), sharing bias data and contributing to the same target distribution. The bias acted on the z-axis defined using the center-of-mass *z*-distance between one central sodium ion and the L177 residues in NavMs. The target distribution was chosen to be flat. The sampling interval was chosen as the entire box length. To keep the sodium ion solute close to the pore, its distance from the pore central axis was restrained to stay below 10 Å by adding a flat-bottom umbrella potential. The rate of change of each AWH bias was initialized by setting the average free-energy error to 20 kj/mol and the diffusion constant to 0.00005 nm^2^/ps. The positions of all C_α_ atoms of the protein were restrained by imposing harmonic potentials with force constants of 1000 kJ mol^-1^ nm^-2^. Convergence was assessed by monitoring the evolution of the free energy profile and target distribution over time. The average potential of mean force profile and associated uncertainties were calculated from a single AWH walker since all the walkers communicate with one another. The average PMF profile was calculated by taking the data from the last 100 ns in intervals of 10 ns and the error bars were estimated by calculating the standard deviation over this dataset. The free energy profiles were then shifted to set the reference to the center of the bilayer and allow a direct comparison with the water density profiles estimated using the channel annotation package (CHAP).

### Drug binding free energy calculation

The free energy of binding of pore blockers lidocaine and flecainide was calculated as described above except for the following parameters. For drugs, the bias acted on the z-axis defined by the center-of-mass *z*-distance between the drug and the E178 residues in NavMs. The sampling interval was limited by harmonic restraints and bounded between 0.8 Å and 3.0 Å. Other simulation details are the same as for the ion permeation free energy calculation described above. Convergence, average profiles and uncertainty estimates were assessed as described above for the ion permeation free energy profiles.

**Table 2:** Summary table of AWH enhanced sampling simulations performed herein.

| System | Production |
| --- | --- |
| NavMs $\alpha$ model - Na<sup>+</sup> | 200 ns X 6 walkers |
| NavMs $\pi$ model - Na<sup>+</sup> | 172 ns X 6 walkers |
| NavMs $\alpha$ model - Lidocaine | 200 ns X 6 walkers |
| NavMs $\pi$ model - Lidocaine | 200 ns X 6 walkers |
| NavMs $\alpha$ model - Flecainide | 200 ns X 6 walkers |
| NavMs $\pi$ model - Flecainide | 200 ns X 6 walkers |
| NavMs $\pi$ model N214L - Na<sup>+</sup> | 200ns X 6 walkers |
| Lidocaine-Membrane | 122ns X 6 walkers |

### Analysis

The water number density and hydration free energy were calculated using Channel annotation package (CHAP) (30, 59). The C-terminal domain was removed from the system for the CHAP analysis. The plots produced show the time-averaged water number densities and hydration free energy. Error bars represent the standard deviation, computed over frames extracted every 1ns over the whole trajectory of the respective systems. Sequence alignments were constructed using CLUSTALW web server with default parameters (60). The orientation of conserved asparagines (Figure 4.D) was evaluated by shifting and reorienting the different structures (PDBIDs 4DXW, 5HK7, 4BGN, 4EKW, 5VB2, 5VB8, 5YUA, 5EK0, 4MVR, 4MVQ, 4MVO, 4MVM, 4MTO, 4MTG, 4MTF, 4MS2, 4MW8, 4MW3, 4MVZ, 4MVU, 4MVS, 3RVZ, 3RW0, 3RVY, 5YUC, 5YUB, 5KLB, 6VWX, 5HVX, 5HVD, 6N4R, 5X0M, 6A90, 5XSY, 6AGF, 6UZ3, 7K18, 6J8I) into a common reference frame: the C_α_ of the conserved Asparagine was placed at the origin (0,0) and the vector connecting the C_α_ atom of the conserved Asparagine and the center of pore was aligned with the x axis. The x and y values in the plot correspond to the x and y components of the vector connecting the C_α_ and the C_γ_ atom of the conserved Asparagine. The interaction energy between sodium ion and individual pore-lining residues was estimated as the sum of coulomb short-range interaction and Lennard-jones short range interaction, using the gmx energy module in GROMACS (57). The files necessary to reproduce the simulations and the analyses reported in this paper are publicly available on OSF: https://osf.io/q4m9w/

## Results

### The full-length NavMs X-ray structure has a dehydrated pore

In order to test the functional annotation of NavMs as a model for gating, we first examined the hydration of the channel pore using MD simulations. The full-length X-ray structure of NavMs (PDB ID 5HVX) was initially proposed to represent the open state, based on its radius at the activation gate (24). However, pore hydration has been considered a prerequisite for conduction in ion channels, and may provide more informative metrics than geometric radius alone (30). Indeed, the capacity of this structure for conduction was recently challenged based on its propensity to dehydrate in MD simulations (36). In our hands, the pore radius of full-length NavMs contracted slightly (∼0.5 A) during equilibration in a lipid bilayer, even with the backbone restrained (Figure S3A-C). During subsequent unrestrained simulations, the pore maintained a stable profile over 400 ns (Figure S3D, S4A,C). Throughout this run, water density was effectively zero near the activation gate (designated -1 nm along the pore axis, Figure 2A). The molecular determinants of this dehydration appeared to be the pore-facing Leu-211 and Ile-215 residues, which created a hydrophobic constriction in this region (Figure 2B) (36). Given the close packing of S6 against the full length of S5 and the S4–S5 linker in this structure (Figure 2C), a local expansion sufficient to hydrate the activation gate does not appear possible, barring substantial rearrangement of the already fully-activated VSD as well as pore domain. Thus, our simulations confirmed that the full-length NavMs structure—termed the α-model, as described in subsequent sections—is unlikely to represent an open state.

**Figure 2:**
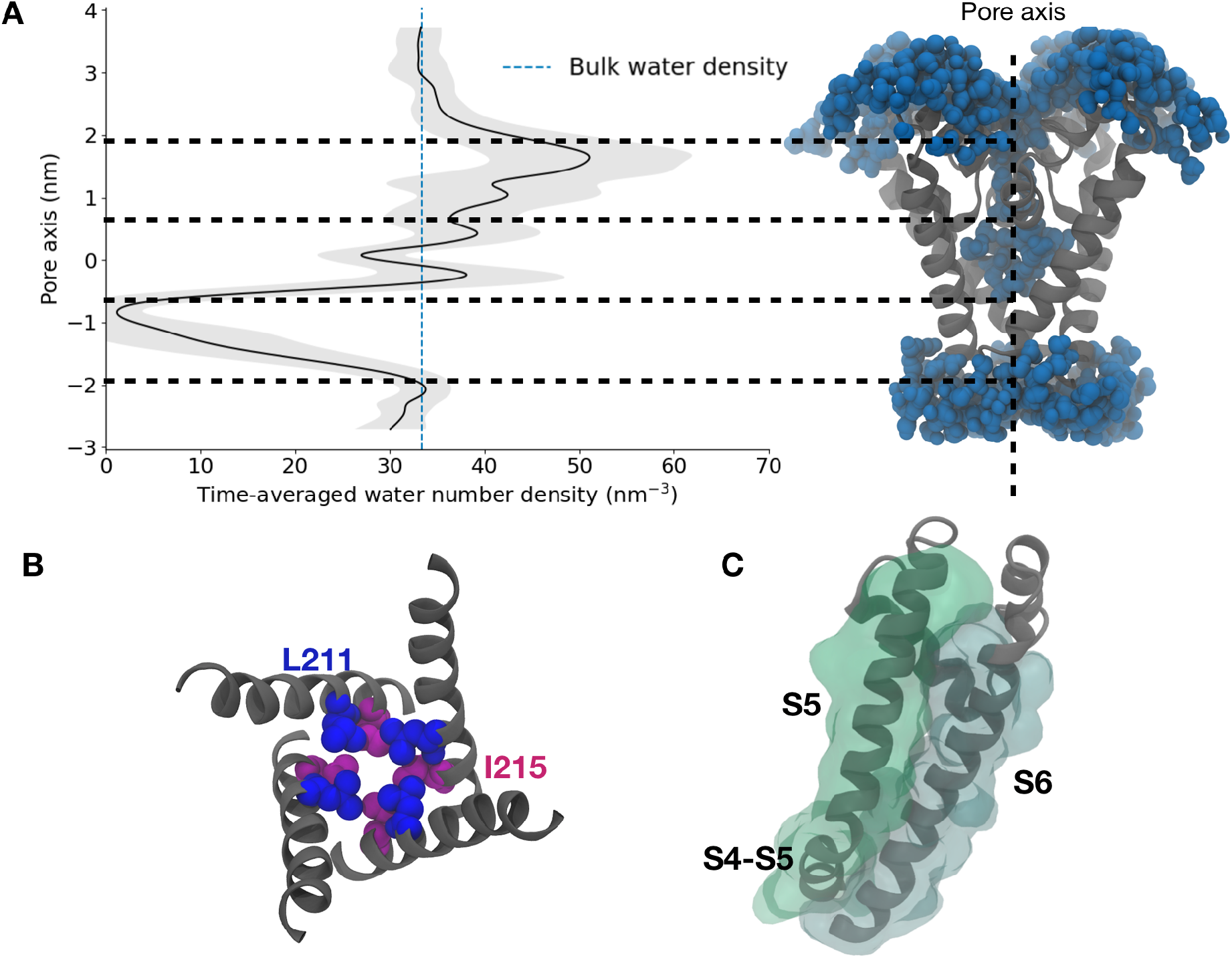
A hydrophobic plug in the full-length NavMs structure (α-model) incompatible with pore hydration. **A** Time-averaged number density of water projected along the pore axis of the α-model, with standard deviation in gray, and bulk-water density as a dashed blue line. Inset at right shows an aligned view of the pore-lining S6 helices in a representative simulation frame, with proximal water molecules in blue. Horizontal dashed lines delineate, from top to bottom, regions corresponding to the selectivity filter, central cavity, and activation gate. The region around -1 nm (activation gate) is clearly dehydrated. **B**. Representative simulation frame for the α-model, showing the S6 helices from the extracellular side. Hydrophobic residues Leu-211 (blue) and Ile-215 (magenta) form a hydrophobic constriction at the activation gate. **C**. Ribbon and semitransparent surface representations showing tight packing of the S5 (green) and S6 helices (gray) from a single NavMs subunit, viewed as in A.

### Transmembrane potential is insufficient to consistently hydrate the pore

Having observed the experimental structure to be nonconducting at 0 mV, we further asked whether the application of a transmembrane potential might stimulate conformational transition to a hydrated, more plausibly open state. Indeed, the introduction of an electrical potential has been shown to alter the surface tension of water at hydrophobic surfaces, and may result in the hydration of transmembrane pores via an electrowetting phenomenon (61, 62). We therefore probed whether electrowetting might also enable hydration of NavMs. To do so, we applied external electric fields resulting in transmembrane potentials of around +750 mV and -750 mV to the so-called α-model, and computed the time-averaged water density during further unrestrained simulations, initially up to 800 ns. Applying external electric fields did indeed increase the pore hydration at the activation gate (Figure 3), but water density remained substantially less than that of the bulk solvent. In fact, the pore repeatedly transitioned between wetted and dewetted states at both +750 mV and - 750 mV (Figure S5). This observation led us to propose that more substantial remodeling is required for the NavMs pore to consistently hydrate.

**Figure 3.**
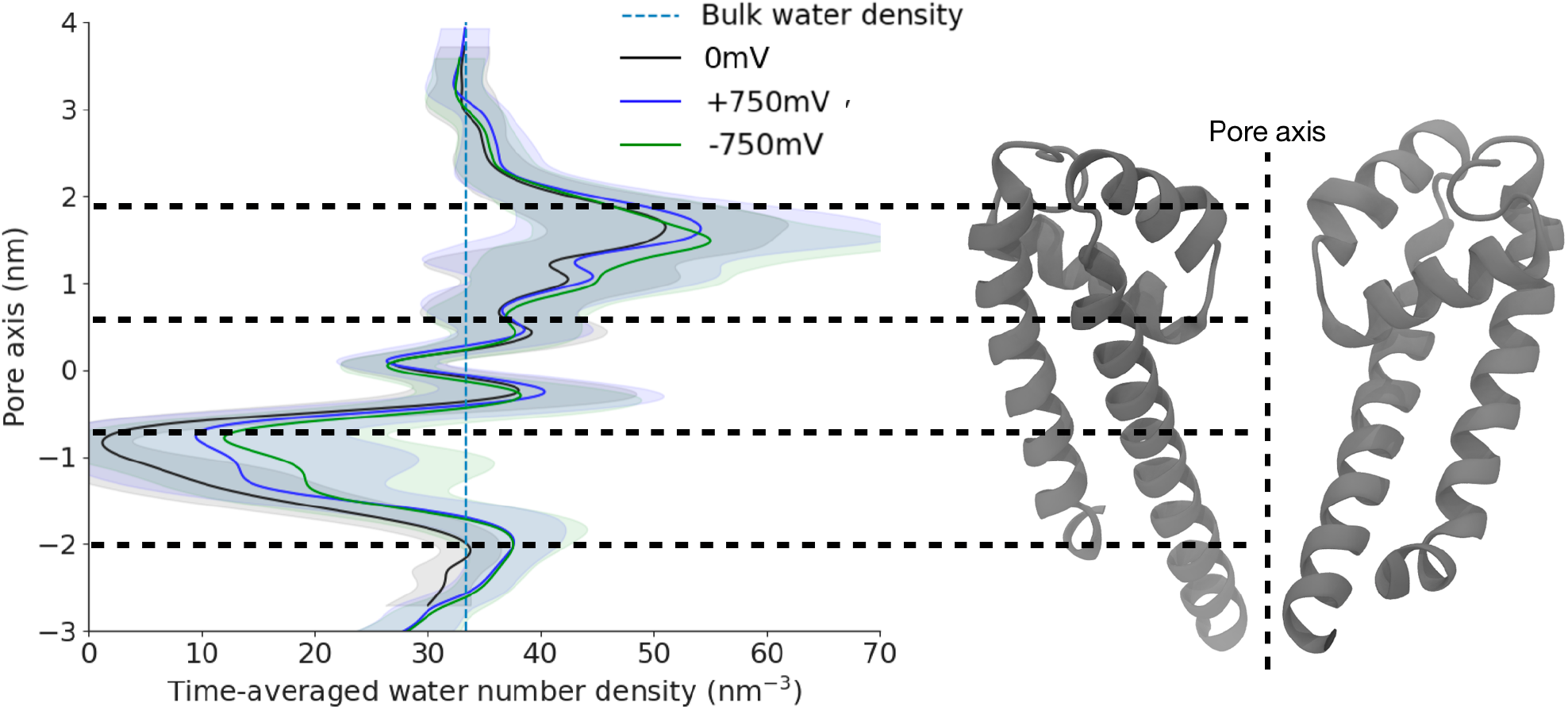
Application of an electric field does not fully hydrate the NavMs α-model pore. Water-number density projected along the pore axis of the α-model, averaged over 800 ns at 0 mV (as in Figure 2, black), +750 mV (depolarized, blue), or -750 mV (hyperpolarized, green), with standard deviations (gray) and bulk-water density (dashed blue). Inset at right shows an aligned view of the pore-lining S5–S6 helices from two subunits. The -1 nm region (activation gate) is only partially hydrated by application of an electric field.

### Formation of a helical defect in S6 and rotation of a conserved asparagine at hyperpolarized potentials

Interestingly, during extended simulations (>8 µs) of NavMs at -750 mV a kink formed spontaneously in the S6 helix of one subunit, with disruption of the backbone-hydrogen bond between residues Val-210 and Phe-214 (Movie S1). Rotation of S6 C-terminal to this kink led the hydrophobic side chains of residues Leu-211 and Ile-215 to reorient away from the activation gate. In their place, an asparagine residue (Asn-212) facing the S4/S5 linker rotated inwards towards the pore (Figure 4A–B). This asparagine is the only hydrophilic residue in the lower S6 segment of NavMs and is conserved across several bacterial and eukaryotic sodium channels, raising the possibility of its importance for channel function (63) (Figure 4.C). Indeed, the equivalent asparagine has also been shown to mediate coupling between the VSD and pore domain in the eukaryotic channels Nav1.4 and Nav1.5 (64) and to come in contact with the inactivation Ile-Phe-Met (IFM) particle of the channels solved in an inactivated state (3, 4, 7, 8).

**Figure 4.**
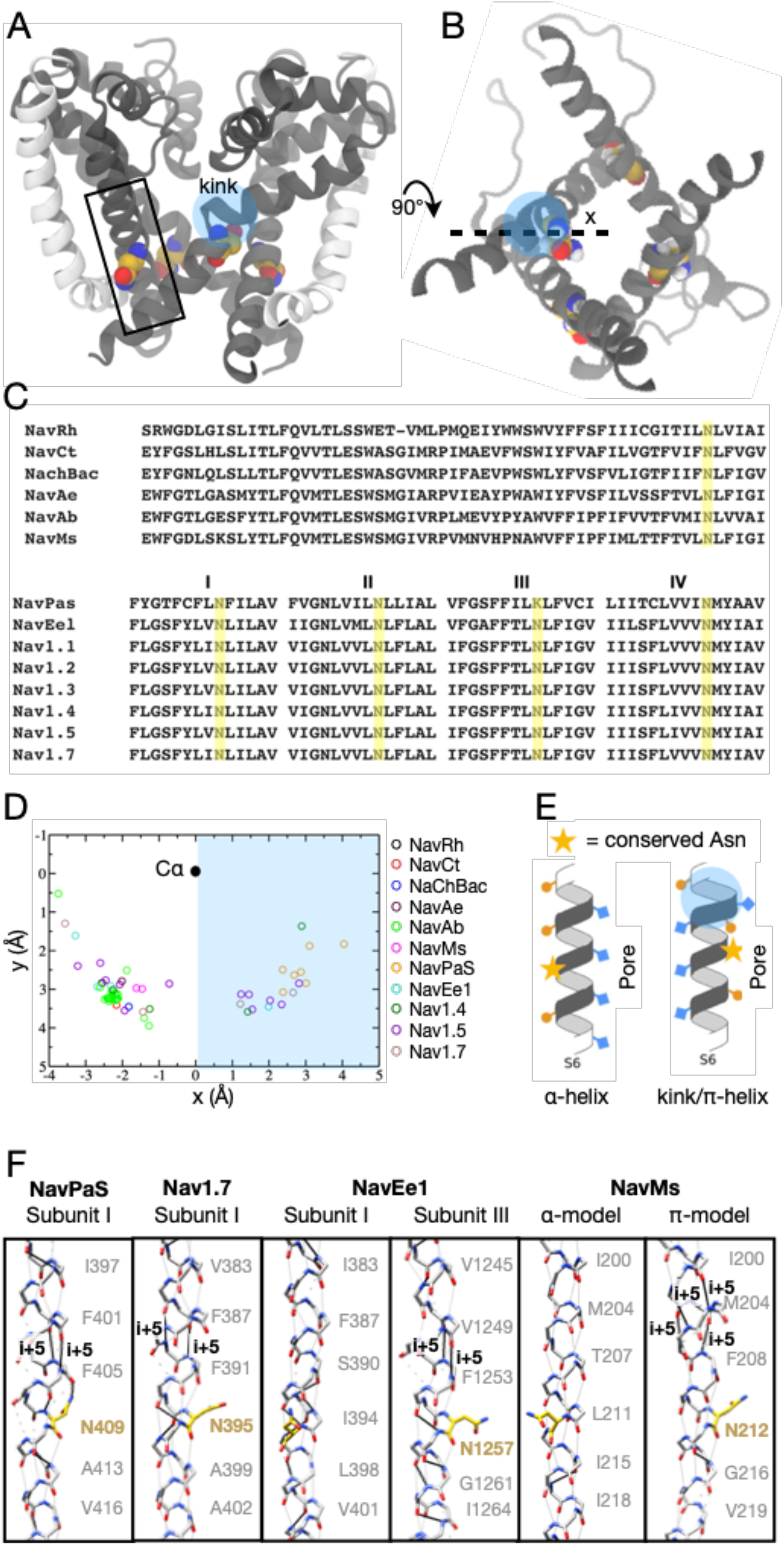
A helix kink and conserved asparagine at the activation gate of NavMs. A) Representative frame from an extended simulation of the α-model at –750 mV, showing the S6 helices (gray) from the membrane plane. A spontaneous kink (shaded blue) is evident in a proximal S6 helix, N-terminal to Asn-212 (yellow). For orientation, S5 helices (white) are shown for the left and right subunits. Solid rectangle indicates zoom region in F. B) Representative frame as in A, viewed from the extracellular side, showing only S6 helices (gray). The kink is evident in the leftmost subunit, with coordinated rotation of Asn-212 towards the channel pore. Dashed line indicates orientation vector used in D, with respect to the leftmost subunit. C) Sequence alignment of S5–S6 regions from six bacterial Nav channels (top), with the conserved asparagine (Asn-212 in NavMs) highlighted in yellow. Alignments of all four pseudohomologous S6 helices in nine eukaryotic Nav channels (bottom) show a similarly conserved asparagine. D) Orientation of the conserved asparagine in eleven sets of experimental structures. For each asparagine, orientation is defined by a projection of Cα–Cγ onto an xy plane perpendicular to the pore (z-) axis, where x is a vector from Cα to the pore center as shown in A. Residues in the shaded-blue region orient towards the channel pore. E) Cartoon depiction of the effect of kink or π-helix formation (shaded blue) on the orientation of a conserved asparagine (yellow star) in S6. In the α-model (left), the asparagine faces S5, away from the pore; in the π-model, it rotates into the pore. F) Homologous regions of S6, as indicated in A with the channel pore at right, in structures of the representative channels NavPaS (PDB ID 6A91), Nav1.7 (PDB ID 6J8I), NavEe1 (PDB ID 5XSY), and the NavMs α-model (PDB ID 5HVX) and π-model. Each structure shows canonical i+4–i (solid gray), disrupted i+4–i (dashed gray), and noncanonical (black) hydrogen bonds; i+5–i interactions, characteristic of π-helices, are labeled. Residue numbers indicate sidechains facing the pore; for clarity, only backbone atoms are shown, except for the conserved asparagine (yellow). Formation of a π-helix is associated with orientation of the asparagine towards the pore (in).

Distortions in S6 have also been observed, where the classical α-helical backbone hydrogen bonding pattern is disrupted and a π-helix formed, along with inward orientation of the asparagine residue, in various subunits of structures of eukaryotic Nav channels (NavEe1, NavPas, Nav1.4, Nav1.5 and Nav1.7, Figure 4D,F). Interestingly, a π-helix transition and asparagine reorientation was recently proposed to be involved in pore opening in the related TRP-channel family (65–67). Based on the spontaneous behavior observed in our simulations, and collective evidence for a conserved gating mechanism, we hypothesized that a S6 distortion and coordinated rotation placing Asn-212 in a pore-facing orientation (Figure 4E,F) might contribute to stabilizing a conductive pore in NavMs.

### π-helix formation in S6 enables pore hydration

To test the plausibility of an open NavMs state containing a π-helix and a pore-facing Asn, we prepared a symmetrized system—termed hereafter the π-model—containing a π-helix and inward-facing asparagine in all four subunits. This model was prepared by introducing a π-helix six positions before the conserved asparagine, using homology modeling to the original structure with gaps at strategic positions in the alignment to shift the sequence by one residue (Figure S1). This realignment resulted in the disruption of backbone hydrogen bonding from the -NH group of residue Thr-207, and rotation of the remainder of S6, leading Asn-212 to reorient towards the pore. During equilibration and production MD simulations, the pore profile was largely stable (pore radius standard deviation within 1 Å, Figure S3.D, S4B,D); notably, Asn-212 remained in a pore facing position, with the unpaired hydrogen bond characteristic of the π-helix motif varying between Thr-207 and Phe-210. Moreover, in contrast to the α-model, the pore equilibrated to ∼3 Å radius and was completely hydrated throughout our unrestrained simulations of the π-model (Figure S4.B, 5A), with water density similar to bulk solvent across the entire pore. Indeed, the free-energy profile for solvation was effectively flat for the π-model, in contrast to barriers up to 3 kcal/mol in our previous α-model simulations (Figure 5B).

**Figure 5:**
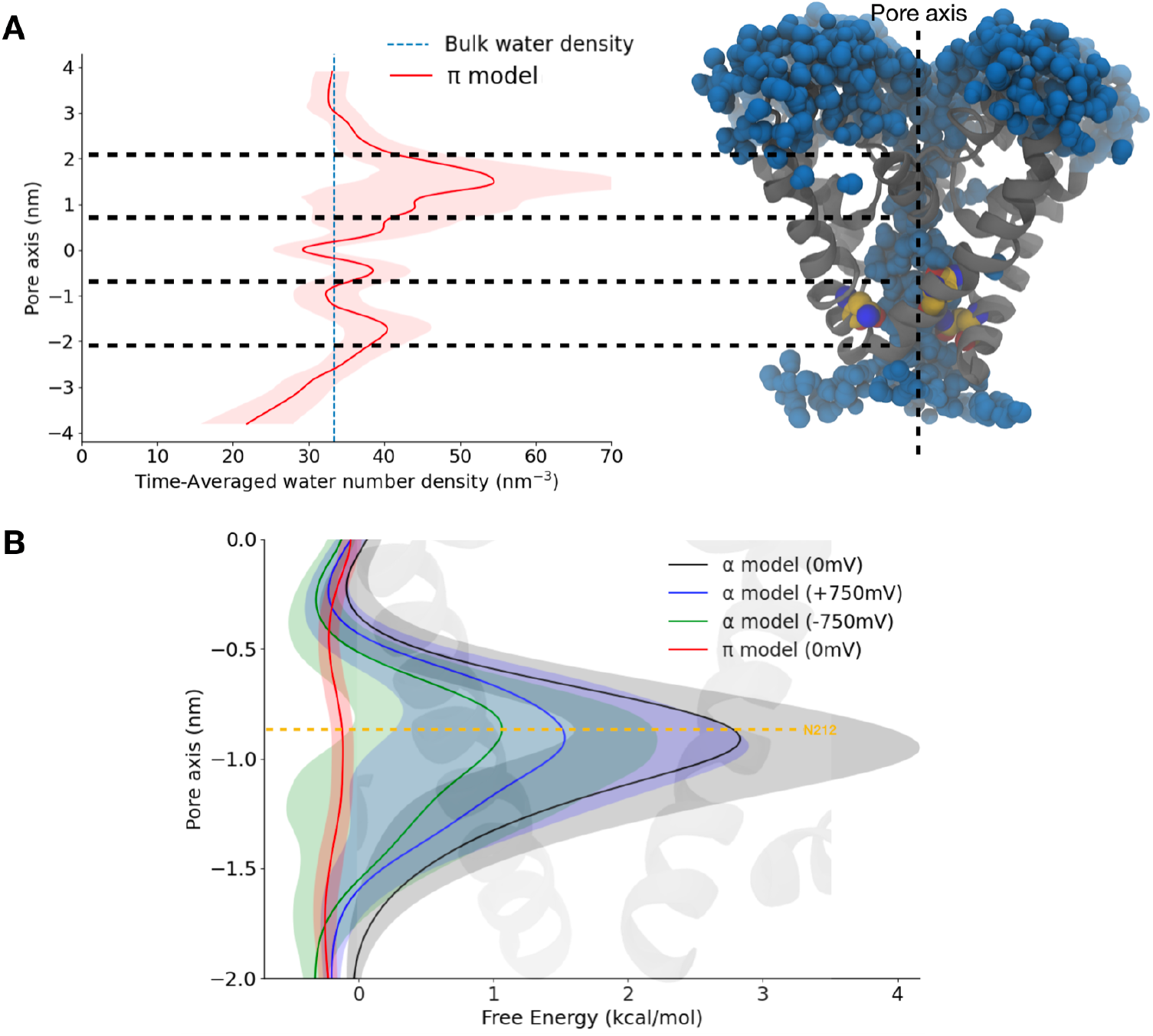
Pore hydration in a NavMs model with an introduced π-helix (π-model). **A**. Time-averaged number-density of water projected along the pore axis of the π-model (red), with standard deviation in transparent representation, and bulk-water density as a dashed blue line. Inset at right shows an aligned view of the pore-lining S6 helices in a representative simulation frame, with proximal water molecules in blue, and the conserved asparagine in yellow. The -1 nm activation gate is clearly hydrated. **B**. Free-energy profiles of pore hydration, derived from water densities, projected along the pore axis for the α-model at 0 mV (black), -750 mV (green), or +750 mV (blue), or for the π-model (0 mV, red). Shaded regions indicate standard deviations; overlaid cartoon shows an aligned view of two opposing S6 helices.

### The π-model conducts Na+ ions

To further probe the conductive properties of our NavMs π-model, we computed the sodium-ion permeation free-energy profiles along the pore axis, using the accelerated weight histogram (AWH) method of enhanced sampling (58). The barrier to sodium conduction was over 25 kcal/mol at the activation gate for the α-model, but was effectively zero throughout the pore for the π-model, indicating that sodium permeates this state readily (Figure 6.A S7, S8). We further tested the contribution of the conserved asparagine to this apparent open state, by running unrestrained simulations with a hydrophobic residue substitute at position 212 in the π-model. Although hydrophobic substitutions (to Cys, Ala or Leu) did not decrease hydration in this model (Figure 6B), sodium ions interacted substantially more favorably with asparagine than with a substituted leucine at position 212 (Figure 6C). Indeed, substitution to Leu increased the barrier to sodium permeation at the gate by over 1 kcal/mol (Figure 6D, S9). These results highlight the importance of monitoring ion as well as water interactions in the channel pore, and indicate a direct role for the conserved Asn in conduction.

**Figure 6:**
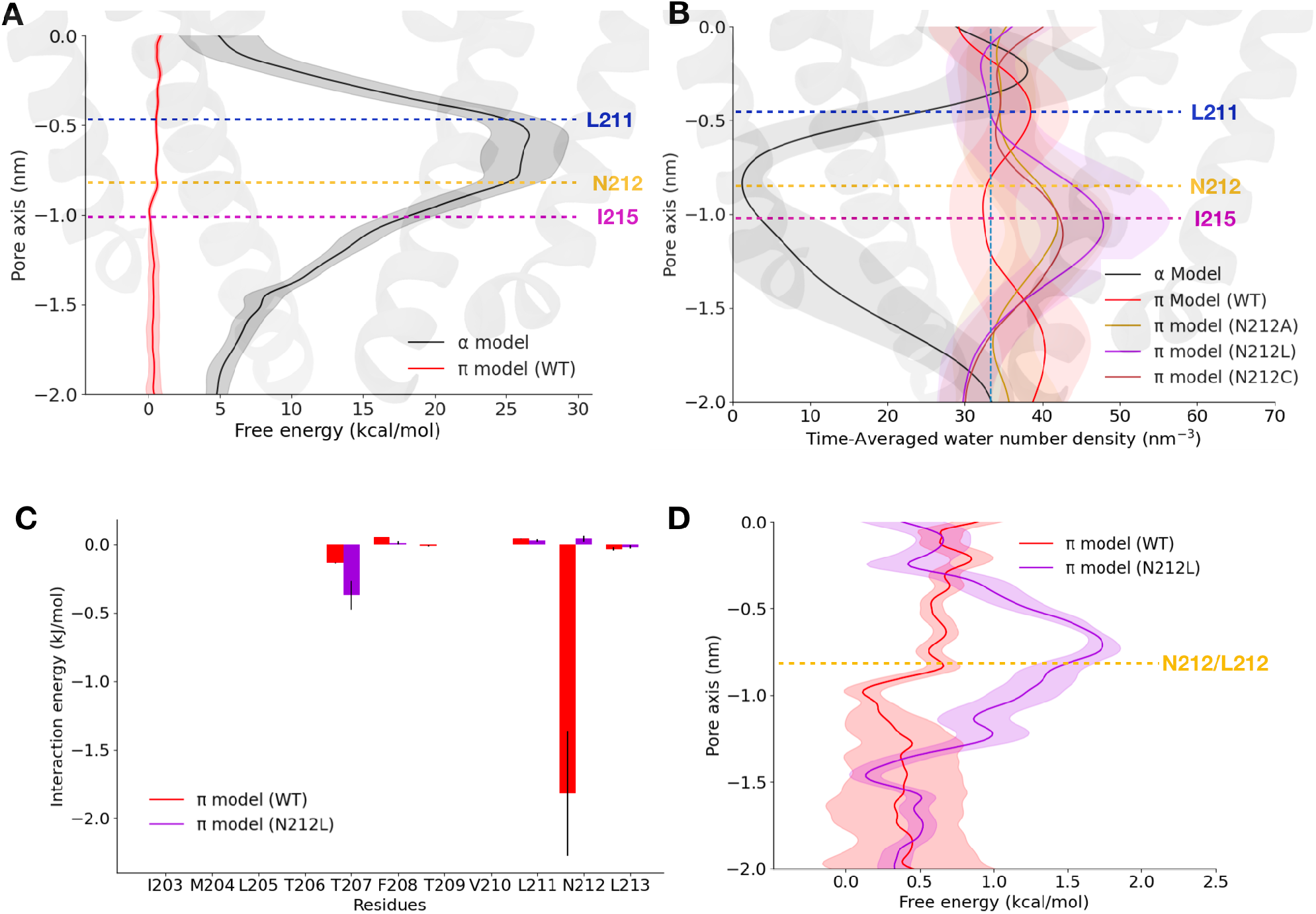
Ion permeation and hydration at the activation gate as a function of π-helix insertion and the presence of the conserved asparagine. **A**. Free-energy profiles for sodium ions in AWH permeation simulations, projected along the pore axis in the α-(black) and π-models (red). Shaded regions indicate standard deviations; overlaid cartoon shows an aligned view of two opposing pore domains. The barrier to sodium permeation is substantial in the α-model, but effectively absent in the π-model. **B**. Time-averaged number-density of water projected along the pore axis for the wild-type α-(black) and π-models (red), and for π-models with mutations at position 212 (alanine, yellow; cysteine, brown; leucine, purple). Hydrophobic substitutions at Asn-212 do not decrease hydration at the activation gate. **C**. Interaction energies with sodium ions for pore-facing residues in the wildtype (red) and N212L (purple) π-models. Favorable interactions are notable at position 212 only when it is asparagine. **D**. Free-energy profiles for sodium ions as in A for the wild-type (red) and N212L (purple) π-models. Leucine substitution moderately raises the barrier to permeation at the activation gate.

### Open-pore blockers bind in the pore cavity of the π-model

As a final test of the functional relevance of our π-model as a NavMs open state, we investigated its capacity for state-dependent drug-binding. According to the guarded receptor hypothesis, open Nav channels must allow open-pore blockers—such as the antiarrhythmic drugs lidocaine and flecainide—to enter the pore cavity from the cytoplasm (Figure S10) (34). We therefore calculated the free energy profiles of binding of lidocaine and flecainide throughout the pore of both the α- and π-models. Similar to Na^+^ permeation free energy profiles, the drug binding free energy profiles for charged lidocaine and flecainide featured large free energy barriers to enter the central cavity via the activation gate in the α-model (barriers > 20 kcal/mol), but were effectively unimpeded in the π-model (Figure 7, S11-14). Interestingly, the binding site of flecainide is similar to the one reported in the α-model of NavAb (11), while it engages in interactions with Asn-212 in the π-model (Figure 7). In contrast, lidocaine appears to interact with Thr-207 in the π-model. Thus, drug-binding as well as ion-permeation and pore-hydration calculations supported the annotation of the NavMs π-model as a putative open state, determined at least in part by the pore orientation of the conserved asparagine.

**Figure 7:**
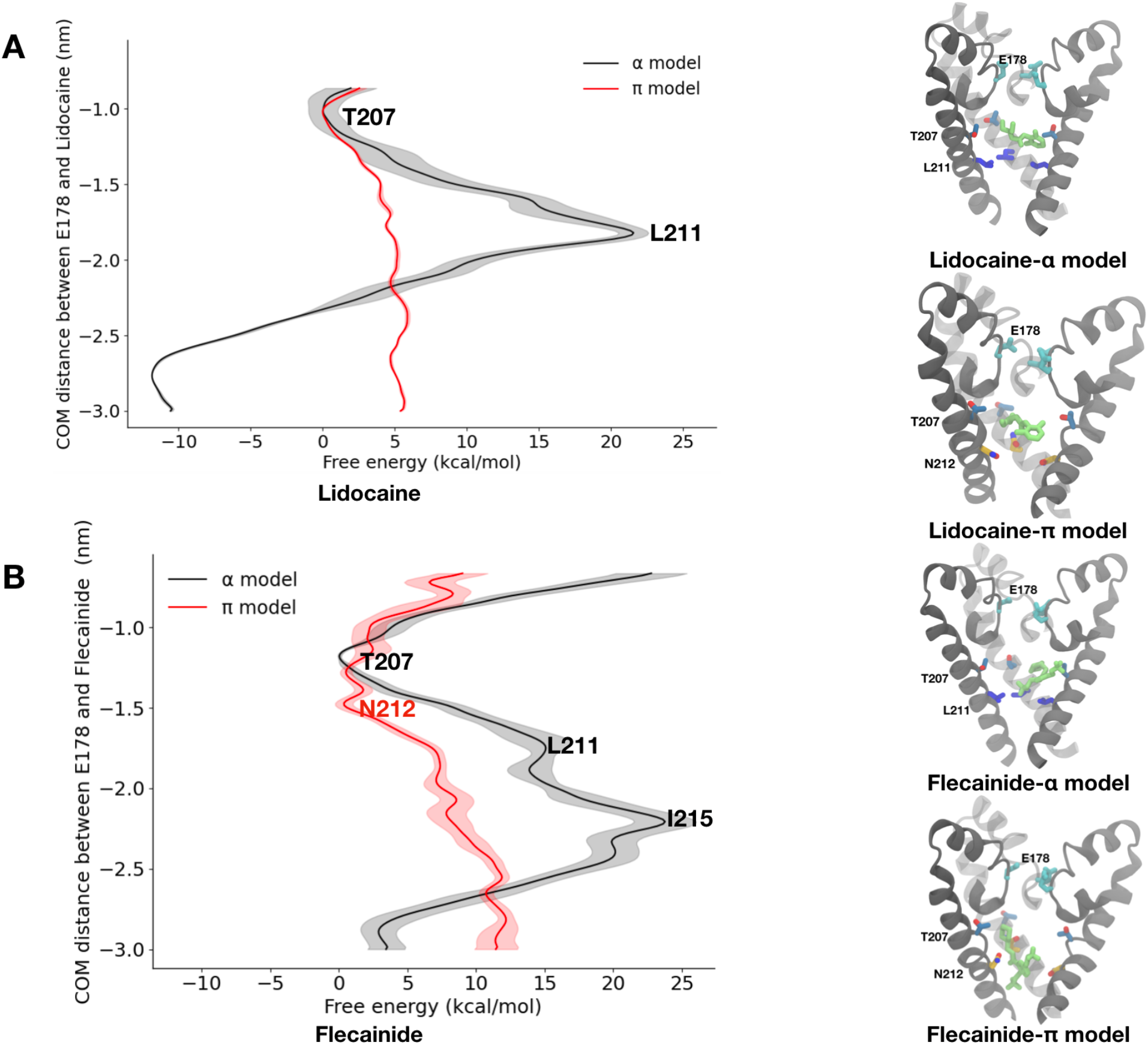
Access of pore blockers to the central cavity in NavMs models. **A**. Free-energy profiles for charged lidocaine in AWH permeation simulations, projected along the pore axis for the α-(black) and π-models (red), with standard deviations shaded. Insets at right show S6 helices in the initial simulation frame for the α-(top) or π-models (bottom), with Asn-212 in yellow and lidocaine in green. **B**. Free-energy profiles for charged flecainide in simulations as in A. Insets at right show S6 helices in the initial frame for the α-(top) or π-models (bottom), with flecainide in green. Barriers to lidocaine or flecainide permeation are substantial at the activation gate in the α-model, but effectively absent in the π-model, enabling either drug to plausibly bind in the central cavity, just above Asn-212 and close to Thr-207 for Lidocaine and at Asn-212 for flecainide

## Discussion

The full-length NavMs structure (α-model) initially appeared to be representative of an open functional state, based on the pore radius estimated at the activation gate (24). MD simulations, however, consistently report that this conformation undergoes dewetting, e.g. hydrophobic gating (36). This dewetting is observed in absence of substantial rearrangements of the pore backbone resulting from contacts between the intracellular domains and the lower parts of the VSD and S4-S5 linker contributing to the stabilization of the backbone helices in a splayed open conformation (68). Instead, such a behavior can be ascribed to subtle side chain rearrangements of hydrophobic residues at the level of the gate (Leu-211 and Ile-215). A conformational change in which the rotation of the C-terminus results from the introduction of a π-helix upstream of Asn-212, on the other hand, leads to full hydration of the pore (π-model). Ion permeation and open pore blocker binding profiles are consistent with the assignment made on the basis of hydration: the original α-model is impermeable to Na^+^ and open pore blockers while the π-model appears to display determinants of an open state. This is consistent with previous MD simulations reporting Na^+^ conduction of α-models only when restraining the protein or inserting mutations that promote pore hydration (36, 69).

An approach we have used to assess the potential of the two models for representing open functional states has been to consider the accessibility of the inner cavity to open pore blocker drugs. According to the guarded receptor hypothesis (GRH), open pore blockers can only access their binding site in the channel when the activation gate is open (figure S10) (34). Channel opening should indeed open the activation gate, opening up an access pathway for the drug to reach its binding site in the pore. Upon inactivation, the drug is trapped in the pore as the activation gate closes. In the inactivated state, drugs cannot access the pore from the intracellular side as the pore is closed (guarding the drug receptor binding site). On top of confirming the validity of the π-model as an open state model, these simulations further revealed the potential molecular determinants of interaction of these drugs and propose a role for Asn-212 in binding flecainide.

An obstructed, non-conductive pore can correspond to a closed functional state (usually coupled to deactivated VSDs) or to an inactivated or pre-open functional state (coupled to activated VSDs). The two latter are difficult to distinguish based on their structural features only. Indeed, structures of another bacterial channel (NavAb) with activated VSD and obstructed pore have been proposed to represent pre-open (10) and inactivated states (15). This functional assignment, however, rests largely on the assumption that the slow-inactivation mechanism is conserved between BacNavs and eukaryotic channels (15, 70), which remains to be firmly established. In addition, the molecular basis for slow-inactivation remains unclear, even in eukaryotic channels, and may involve several functional and structural states (70, 71). Toxin binding and mutations in the outer pore region, selectivity filter, pore helix, and S6 have all been shown to modify the slow inactivation phenotype of Navs (72–84). Whether the α-model of NavMs is then likely to represent one of the inactivated states is a possibility that remains to be explicitly tested.

Our extensive simulations of the α-model of NavMs under depolarized potentials showed a spontaneous conformational change, namely a disruption the α-helical h-bonding pattern and the formation of a defect reminiscent of a π-helix formation, and the consequent rotation of Leu-211 and Ile-215 away from the pore lumen coupled to the rotation of Asn-212 into it. How might this rotation be energetically allowed? At first glance, the introduction of a h-bond defect in an α-helix might seem prohibited. Nevertheless, structures of both Nav and TRP channels containing unpaired h-bonds involved in π-helices have been resolved, suggesting such a conformational change is energetically accessible (Figure 4F). In addition, our simulations showed that a similar change might be triggered by high-enough transmembrane potentials. Energetic stabilization from protein-protein contacts, or protein-solvent might compensate the energetic cost associated with disrupting a canonical α-helical structure (38, 85). Our simulations suggest that pore hydration, presumably due to the application of a prolonged high transmembrane potential (Figure S6) has presumably favored the reorientation of the sole hydrophilic residue into the hydrated pore.

Asn-212 has previously attracted attention as the only hydrophilic residue in an otherwise hydrophobic region (86). Furthermore, this residue has an outstanding conservation in Bac-Navs and eukaryotic Navs (63), and is also conserved in TRP channels, which are only distantly phylogenetically related. In TRPV1, pore wetting occurred as a consequence of Asparagine residues localizing in a pore-facing conformation (65), thus demonstrating the interplay between hydration and Asn orientation. Nevertheless, contrary to our expectations, our in-silico mutagenesis study showed that a substitution of hydrophilic Asn-212 by hydrophobic residues did not affect pore hydration, demonstrating that the physicochemical properties of the residue at this position are not a crucial determinant of pore hydration. Instead, it appears that it is rather the removal of Leu-211 and Ile-215 from the pore constriction that results from the helix rotation that are determinant for the transition from an obstructed dehydrated state to a hydrated conductive one.

While the importance of the Asn residue is clearly substantiated by its conservation throughout evolution, its specific role in our model thus remains quite mysterious. One aspect that appears important is that it plays a role in the ion permeation process, by interacting favorably with permeating Na^+^ ions. Its substitution by a hydrophobic residue indeed raises the permeation free energy barrier, possibly playing a role in determining single channel conductance. In addition, the conserved Asn may play a role in inactivation, though the structural basis for this phenomenon remains largely unknown. The mutation of this Asn to Asp in NaChBac leads to enhanced inactivation, while the mutation to any other residue is non-functional (86). In eukaryotic channels, the conserved Asn itself has been implicated in inactivation of Nav1.4 and Nav1.2a (79, 87), and a direct interaction between the Asn on DIV-S6 (facing away from the pore lumen) and the IFM particle responsible for fast inactivation is found in structures of eukaryotic Nav channels solved in inactivated states (3, 4, 7, 8). Of particular note, in all these structures, the S6 helix of DIV is systematically α-helical while the S6 of the other domains may feature h-bonding defects and formation of π-helix segments coupled to the orientation of the conserved Asn into the pore domain (Figure 4.D,F). Thus, while we are so far unable to pinpoint the role of this Asn residue, a set of clues point towards its implication in inactivation in Nav channels.

Taken together, our study also highlights the relevance of molecular modeling and MD simulations for improving our understanding of structure/function relationship in ion channels. Nevertheless, one should be mindful of the limitations of the technique, and in particular of the imprecisions of the interaction model (force field) at the basis of the algorithm. Water models may be of specific concern in this study, given that their ability to reproduce the behavior of confined water droplets or single water molecules has been questioned. Here, we have used the CHARMM force field with the TIP3P water model, a common choice when simulating membrane proteins. We note that previous work on very similar systems had probed the effect of changing water models on the hydration of the NavMs pore. Simulations carried out with the AMBER force field had revealed no major discrepancies (36), giving us confidence that these findings are robust with respect to the choice of force field.

## Conclusion

In conclusion, this work confirmed that the full-length NavMs structure reflects a non-conductive state, which is possibly reflective of an inactivated channel. In addition, based on spontaneous conformational change and scrutinizing experimental structures of eukaryotic Nav channels, we proposed a structural change that leads to the pore assuming a conductive state: introducing a π-helix results in the rotation of S6 in a manner compatible with pore hydration. This conformational change is coupled to the orientation of a conserved Asn residue into the pore. Since this residue is conserved throughout Nav channels, we hypothesize that such an opening model may apply to eukaryotic channels.

## Acknowledgements

We acknowledge SciLifeLab and the Swedish Research Council (VR 2018-04905, 2017-04641 and 2019-02433) and Swedish e-science Research Center (SeRC) for funding. The MD simulations were performed on resources provided by the Swedish National Infrastructure for Computing (SNIC) on Beskow at the PDC Centre for High Performance Computing (PDC-HPC) and by PRACE on Piz-Daint at the Swiss national supercomputing center (CSCS).

## Author contributions

K.C, M.K, S.M and L.D designed the research. K.C, M.K and S.M performed the experiments and analyzed the results. All the authors interpreted the data. K.C, R.H and L.D drafted the manuscript.

