## Supplementary figures for "An open state of a voltage-gated sodium channel involving a π-helix and conserved pore-facing Asparagine"

```
                                                         207   212                   234
Template EWFGDLSKSLYTLFQVMTLESWSMGIVRPVMNVHPNAWVFFIPFIMLT-TFTVLNLFIGIIVDAMAITKEQEEEAKT
Target   EWFGDLSKSLYTLFQVMTLESWSMGIVRPVMNVHPNAWVFFIPFIMLTTFTVLNLFIGIIVDAMAITKEQEEEAKT-
                                                         207  212                   234
```

**Figure S1**: Sequence alignment of the S6 helix used as an input to Modeller to build the π-model. The gap is highlighted in blue and the conserved Asparagine in yellow.

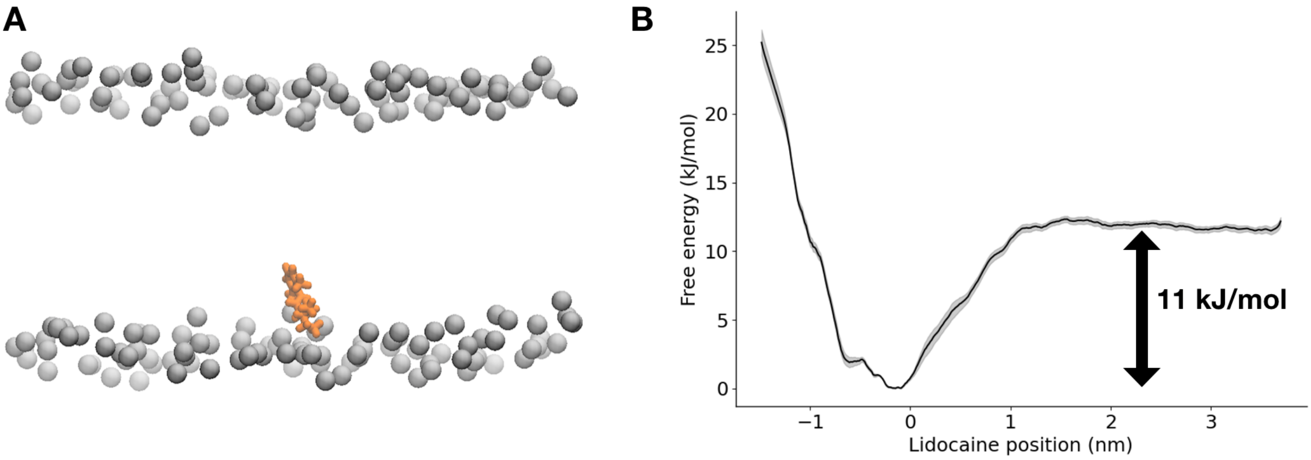


**Figure S2. A.** Lidocaine localization in a POPC membrane. **B.** The lidocaine parameters were checked by calculating the free energy for water to membrane transition along the membrane normal. The free energy difference (indicated by the double headed arrow) for water to membrane transition is around -11 kJ/mol.

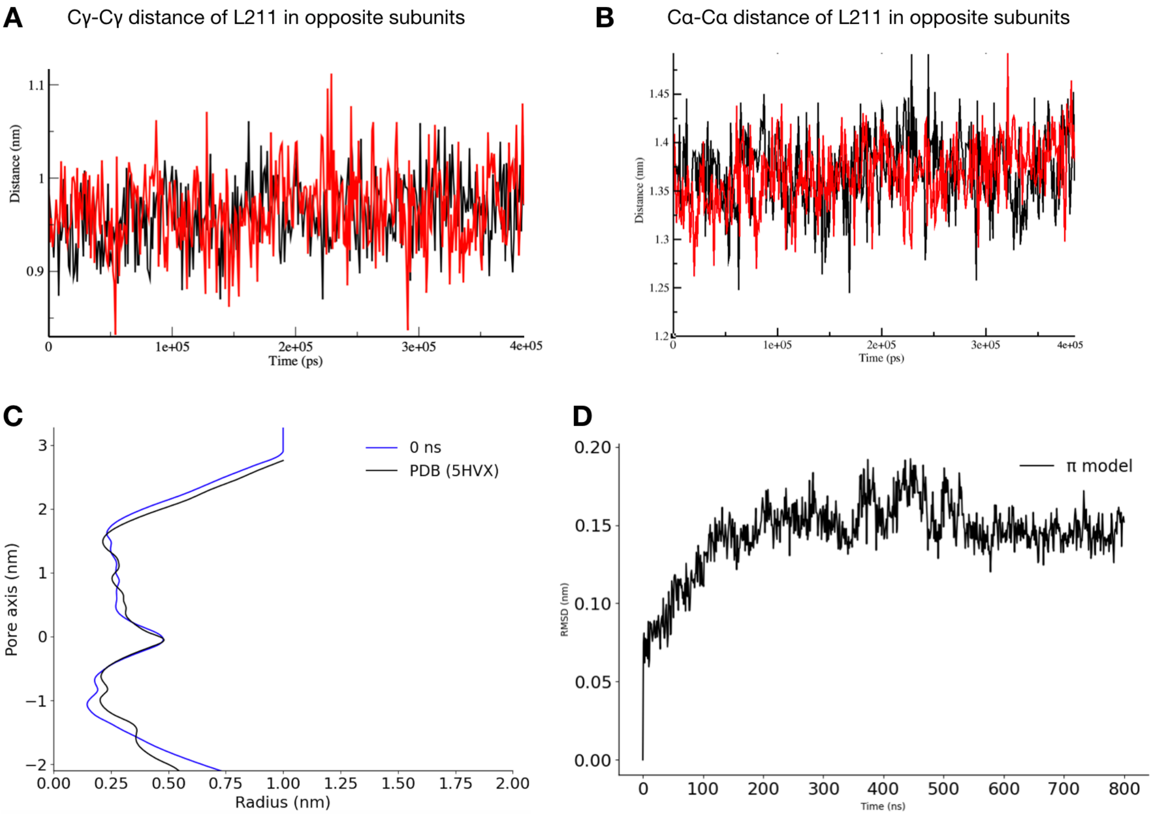


**Figure S3. A.** Leu-211 opposite subunits $C_\gamma$-$C_\gamma$ distance in the α model. The two distances are shown in black and red. **B.** Leu-211 opposite subunits $C_\alpha$-$C_\alpha$ distance in the α model. The two distances are shown in black and red. **C.** Pore radius profile of NavMs PDB structure (PDB ID 5HVX - black) and α model after 48 ns of restrained equilibration. **D.** Root mean squared deviation (rmsd) of the backbone atoms of the four S6 helices of the π model.

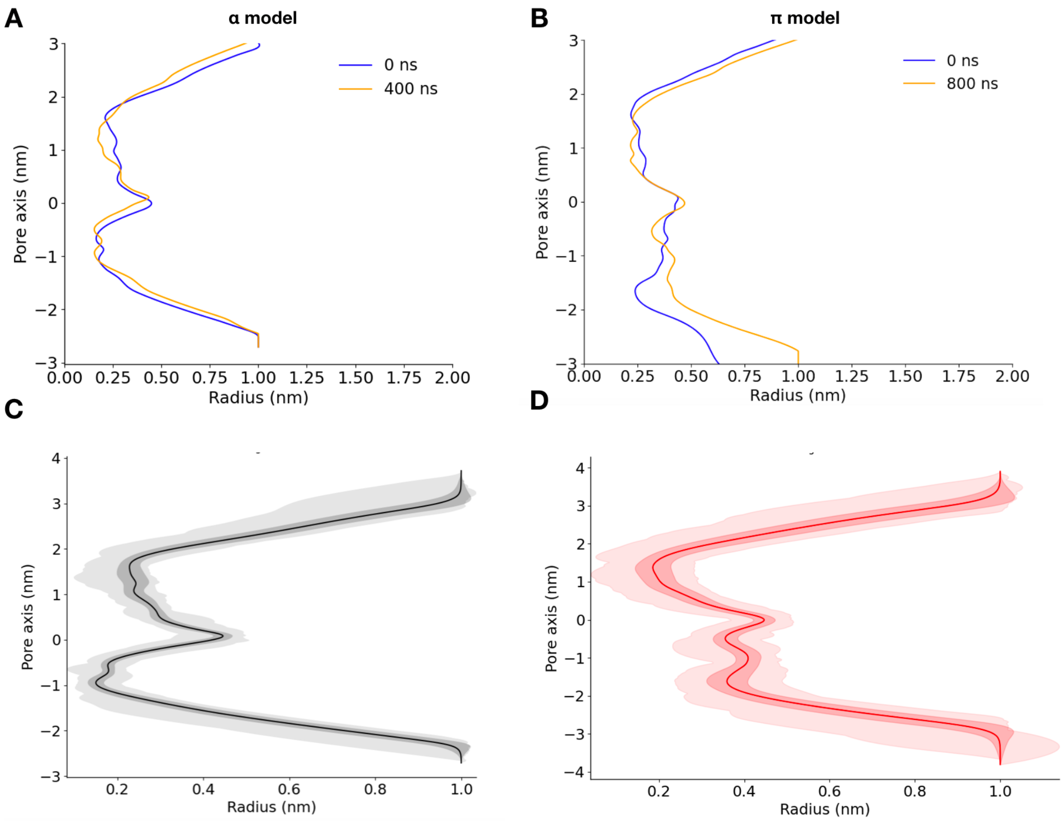


**Figure S4: A.** Pore radius profile of the α model at the start (0 ns) and end (400 ns) of the equilibrium simulation **B.** Pore radius profile of π model at the start (0 ns) and end (800 ns) of the equilibrium simulation **C.** Time-averaged pore radius profile for the α model **D.** Time-averaged pore radius profile for the π model. Dark shaded region show the standard error on the mean, light shaded region the extreme values.

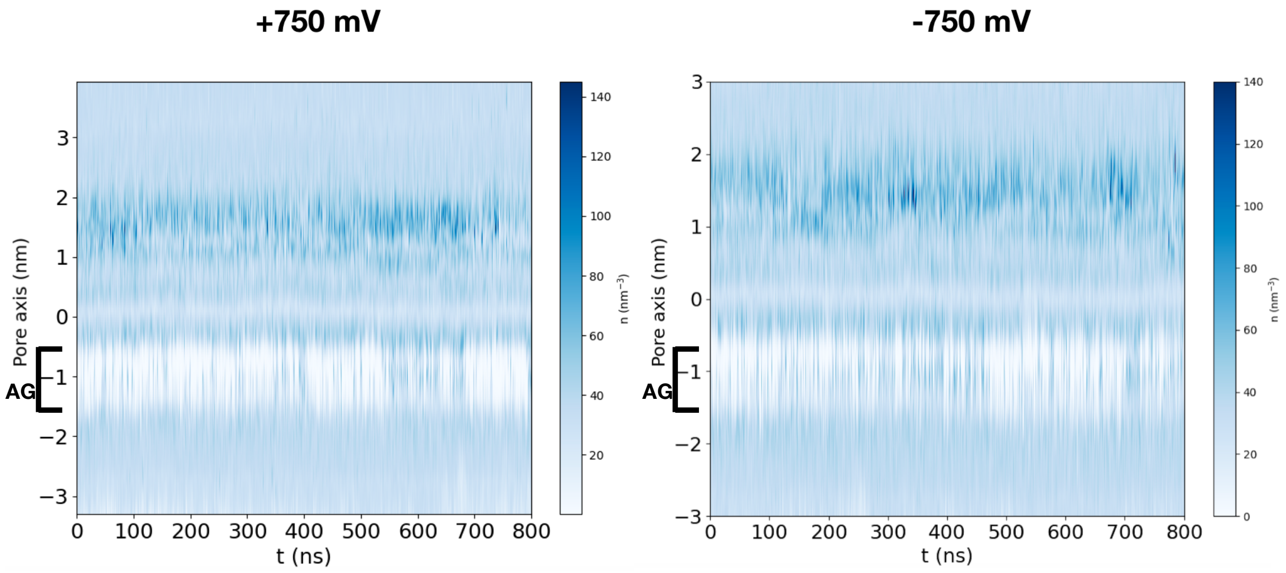


**Figure S5: A.** Water number density profile over time along the central pore axis at +750 mV. The labelled activation gate (AG) is transiently hydrated **B.** Water number density profile over time along the central pore axis at -750 mV. The activation gate is transiently hydrated

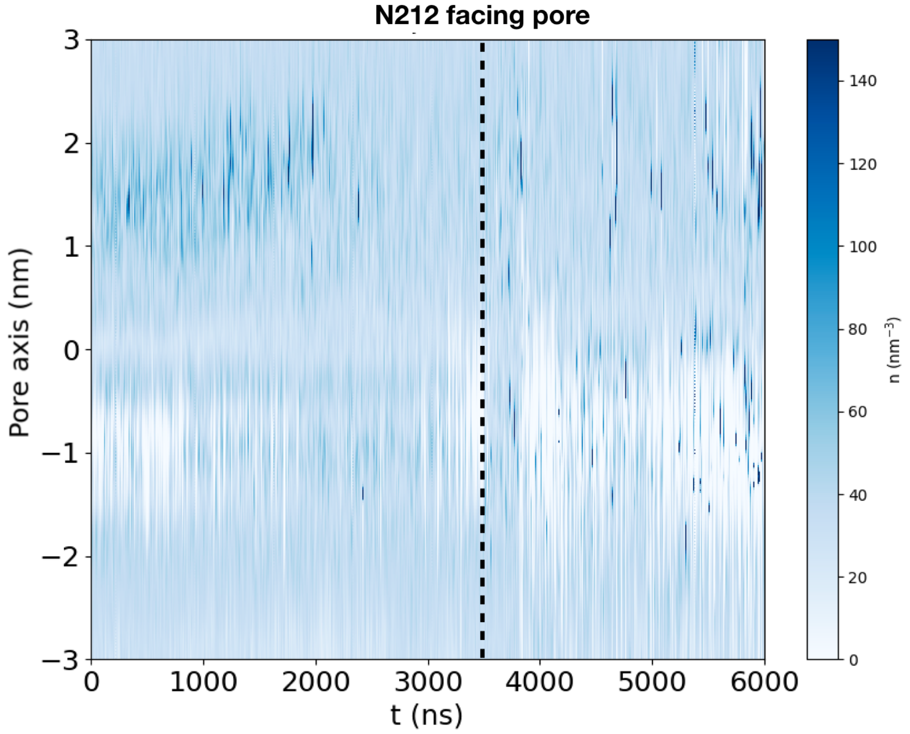


**Figure S6:** Water number density profile over time along the central pore axis at -750 mV. The activation gate is transiently hydrated, preceding the kinking of one of the S6 helices and subsequent reorientation of the conserved Asn into a pore-facing position (black dotted line).

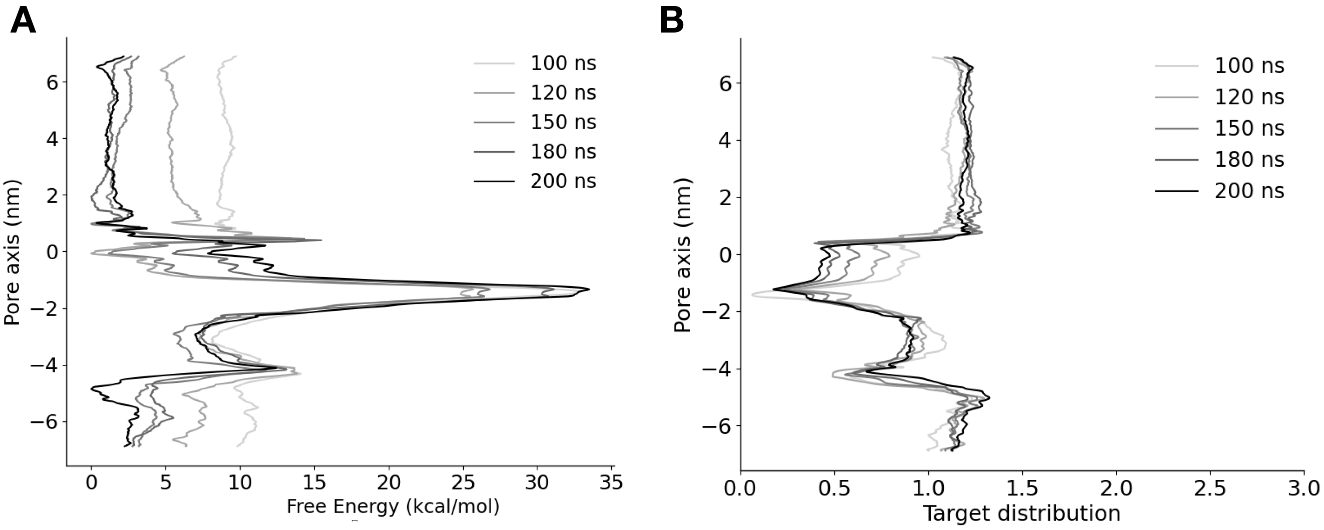


**Figure S7: A.** Convergence of free energy profile of sodium ion permeation in the NavMs $\alpha$-model. The free energy was calculated across 6 walkers sharing the bias. **B.** Target distribution at different times.

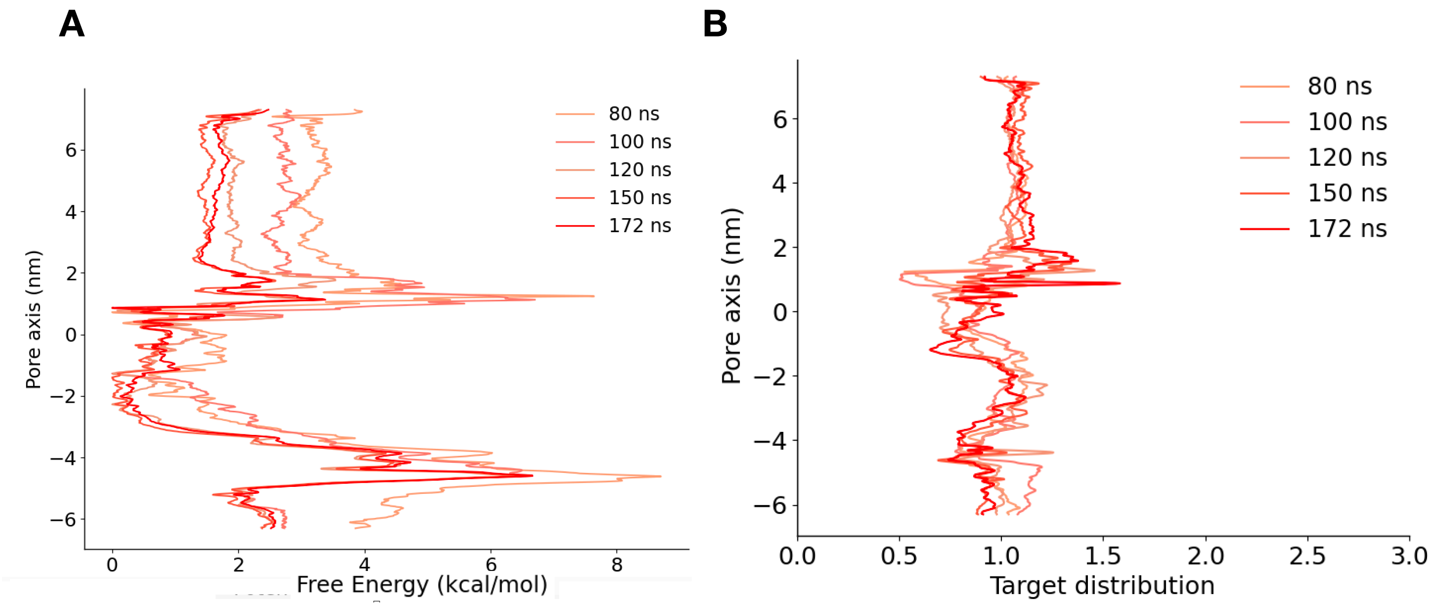


**Figure S8: A.** Convergence of free energy profile of sodium ion permeation in the NavMs $\pi$-model. The free energy was calculated across 6 walkers sharing the bias. **B.** Target distribution at different times

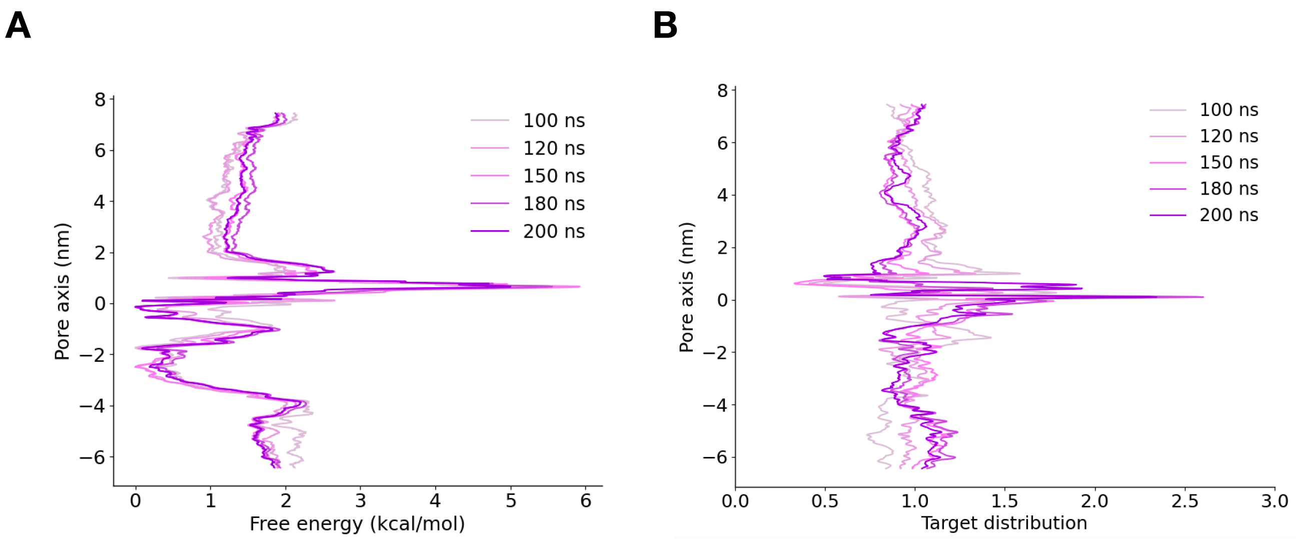


**Figure S9: A.** Convergence of free energy profile of sodium ion permeation in the NavMs $\pi$-model mutant N212L. The free energy was calculated across 6 walkers sharing the bias. **B.** Target distribution at different times

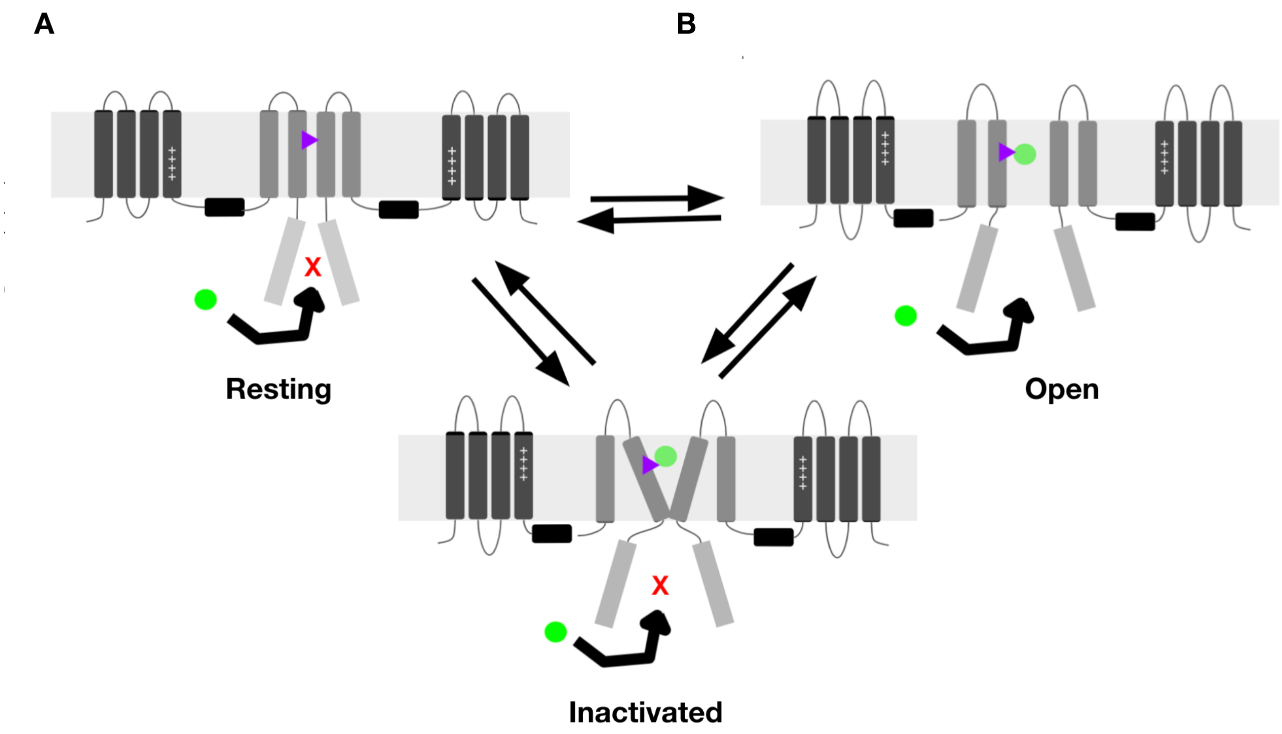


**Figure S10:** Guarded receptor hypothesis. In the resting and inactivated states, the pore is closed. This prevents the drug (green circle) from accessing its binding site (shown as purple triangle) inside the pore. In the open state, the pore is open which allows the drugs to access its binding site without any hindrance.

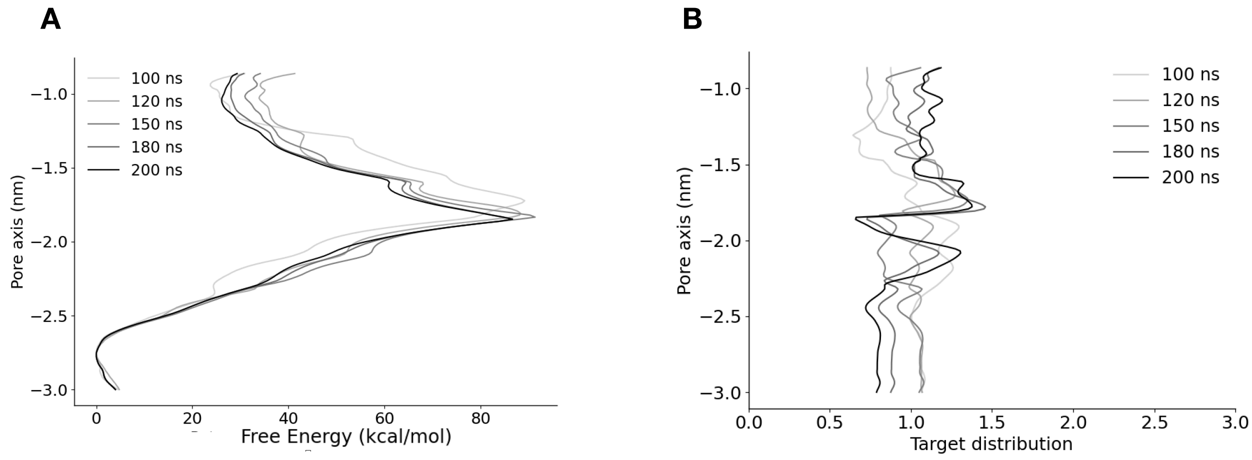


**Figure S11: A.** Convergence of free energy profile of lidocaine permeation in the NavMs $\alpha$-model. The free energy was calculated across 6 walkers sharing the bias. **B.** Target distribution at different times

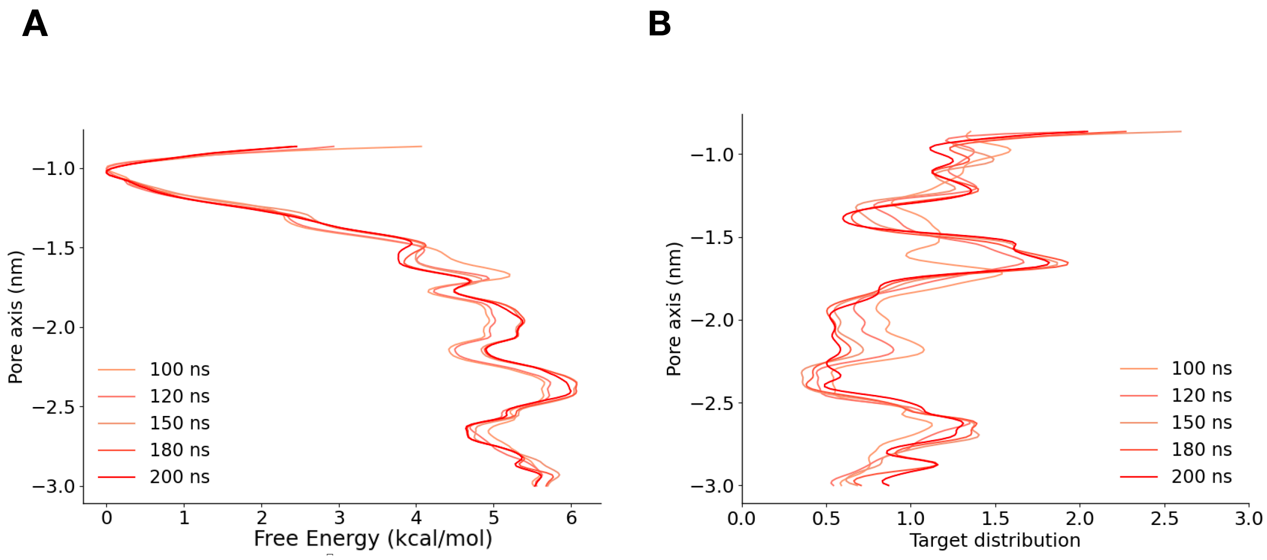


**Figure S12: A.** Convergence of free energy profile of lidocaine permeation in the NavMs $\pi$-model. The free energy was calculated across 6 walkers sharing the bias. **B.** Target distribution at different times

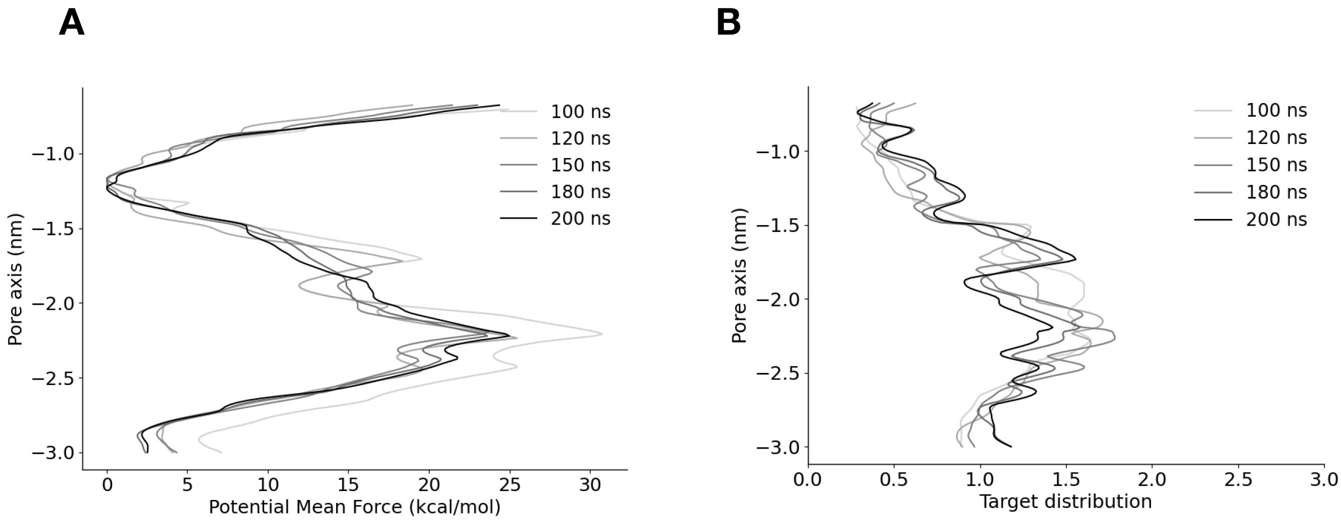


**Figure S13: A.** Convergence of free energy profile of flecainide permeation in the NavMs $\alpha$-model. The free energy was calculated across 6 walkers sharing the bias. **B.** Target distribution at different times

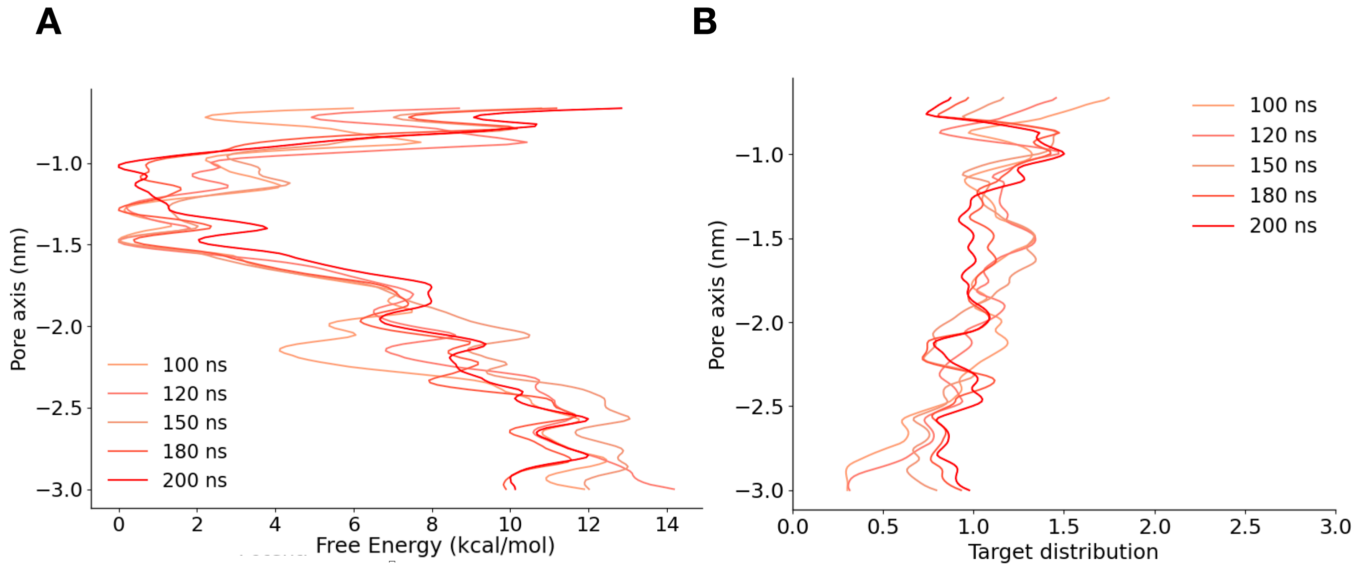


**Figure S14: A.** Convergence of free energy profile of flecainide permeation in the NavMs $\pi$-model. The free energy was calculated across 6 walkers sharing the bias. **B.** Target distribution at different times

**Movie S1** NavMs undergoes a conformational transition when placed under a large hyperpolarizing transmembrane potential. A kink forms spontaneously in the S6 helix (grey ribbon) of one subunit. Rotation of S6 C-terminal to this kink leads Asn-212 (yellow) to rotate inwards towards the pore.
